# Custom-made design of metabolite composition in *N. benthamiana* leaves using CRISPR activators

**DOI:** 10.1101/2021.07.12.452005

**Authors:** S Selma, N Sanmartín, A Espinosa-Ruiz, S Gianoglio, MP Lopez-Gresa, M Vázquez-Vilar, V Flors, A Granell, D Orzaez

## Abstract

Transcriptional regulators based on CRISPR architecture expand our ability of reprogramming endogenous gene expression in plants. One of their potential applications is the customization of plant metabolome through the activation of selected enzymes in a given metabolic pathway. Using the previously described multiplexable CRISPR activator dCasEV2.1, we assayed the selective enrichment in *Nicotiana benthamiana* leaves of four different flavonoids, namely naringenin, eriodictyol, kaempferol and quercetin. After careful selection of target genes and guide RNAs combinations, we created successful activation programs for each of the four metabolites, each program activating between three and seven genes, and with individual gene activation levels ranging from 4- to 1500-fold. Metabolic analysis of the flavonoid profiles of each multigene activation program showed a sharp and selective enrichment of the intended metabolites and their glycosylated derivatives. Remarkably, principal component analysis of untargeted metabolic profiles clearly separated samples according to their activation treatment, and hierarchical clustering separated the samples in five groups, corresponding to the expected four highly enriched metabolite groups, plus an un-activated control. These results demonstrate that dCasEV2.1 is a powerful tool for re-routing metabolic fluxes towards the accumulation of metabolites of interest, opening the door for custom-made design of metabolic contents in plants.

## Introduction

Traditionally, plant breeding has contributed to the generation of crop varieties adapted to changing external conditions and to consumers’ demands by selecting favourable genetic variants in a species’ genomic pool, or by introducing new genetic traits through transgenesis or mutation by CRISPR (Jisha et al. 2015; Maioli et al. 2020). In recent years, the need for new adaptations has grown exponentially, fostered by climate change and human population dynamics, and the pressure to explore new breeding strategies has increased (Godwin et al. 2019). In this context, new breeding concepts inspired in Synthetic Biology propose the development of new programmable traits where physiological outputs (e.g., a developmental phase transition, or the activation of a metabolic pathway) occur as a response to endogenous or external inputs (e.g., a chemical cue, or an electromagnetic signal) perceived by synthetic sensors, and operated by engineered genetic operators (e.g., logic gates, toggle switches, oscillators, etc) (McKenzie et al. 1998; Ochoa-Fernandez et al. 2016; Bernabé-Orts et al. 2020). To produce the desired physiological output, operators need to be transcriptionally connected to a selected group of final actuators (e.g., a group of enzymes), that ultimately generate the designed phenotypic changes. Natural gene circuits have evolved intricated regulatory cascades of transcription factors (TFs) that connect operators with downstream actuators in a concerted manner, jointly generating a consistent physiological response. Among the many challenges facing Plant Synthetic Biology, a key one is to acquire the ability to customize the connections between synthetic operators and endogenous actuators in ways that are different to those designed by evolution but convenient for agriculture. Examples of new “synthetic” connections are the modification of flowering time (Papikian et al. 2019), the activation of an anticipated response to a forecasted biotic/abiotic stress (Chen et al. 2020), or the customization of metabolic composition (Llorente et al. 2020). Natural transcription factors, which are often used as connection hubs in traditional genetic engineering approaches (Xie et al. 2006; Naing et al. 2017), have a limited capacity for circuit customization due to their hardwired DNA binding specificities, which impede free selection of the downstream genes to be regulated. Recently, a new type of programmable transcriptional regulators (PTRs) based in CRISPR/Cas has emerged that allow easy customization of both DNA binding and transcriptional regulatory activities. The nuclease-inactivated CRISPR/Cas9 (dCas9) (Maeder et al. 2013) architecture enables the combination of autonomous activation domains (ADs) with the DNA-binding specificities of Cas9, which can be programmed through a 20 nucleotide-long guide RNA with minimum engineering efforts. In plants, several strategies to build potent these Programmable Transcriptional Activators (PTA) based in dCas9 have been described (Park et al. 2017; Lowder et al. 2018; Li et al. 2019; Pan et al. 2021), showing efficient activation of target endogenous genes. The two main advantages of dCas9-PTA are (i) their high accuracy, reaching single-gene specificity levels as shown recently in transcriptomic analysis showing negligible off-target activation (Li et al. 2017); and (ii) their amenability for multiplexing (Vazquez-Vilar et al. 2016; Lowder et al. 2017). This latter feature enables the concerted activation of multiple actuators simultaneously, by simply loading the cell with several gRNAs, each one targeting a different promoter or a different position within a given promoter. The practical implications of multiplexing PTRs are widespread, from the design of synthetic regulatory cascades to the re-routing of endogenous metabolic fluxes. However, the capacities of PTRs in producing new-to-nature phenotypes are just starting to be explored, and no examples exist yet where PTRs are applied to re-route biosynthetic fluxes. Mastering the regulation of metabolic pathways would open the way for the customization of plant composition, a possibility with many implications in food and feed design, as well as in the development of plant biofactories.

The phenylpropanoid pathway is an alluring bioengineering target due to its pharmaceutical and industrial interest (Neelam et al. 2020). Besides, it is a highly branched pathway in plant secondary metabolism, offering interesting opportunities for biotechnological regulation and fit for the purpose of testing new technological approaches. The pathway can be divided into different parts (see Figure 1). The “general” section of the pathway generates cinnamic acid, coumaric acid and 4-coumaroyl-CoA, the basic backbone products derive from phenylalanine, thus providing the core structures for the biosynthesis of all flavonoids as well as for other phenylpropanoid branches like the lignin pathway; a second group of reactions leads to the condensation and subsequent cyclization of the core structure, generating the first flavanones of the pathway; finally, flavanones will be the substrate for multiple reactions that originate the distinct subgroups of the flavonoid pathway, including flavonols, anthocyanins, isoflavonoids, etc (Nabavi et al. 2020). In plants, the main flavanones are naringenin, eriodictyol and hesperitin, which can also be found in their glycosylated forms and whose distribution changes between species and tissues. For example, in grapefruit and tomato naringenin glycosyl-conjugated compounds are the predominant flavanones present in the fruits, while in some species of citrus, like mandarin or lime, hesperitin glycosyl-conjugates are the most abundant flavanones (Khan et al. 2014). This variation is due to the different combination of gene expression patterns in the upstream part of the pathway. Similarly, the differential expression of downstream enzymes governs the predominant accumulation of flavonols (e.g. kaempferol, quercetin) or anthocyanins (e.g. delphinidin, pelargonidin), shaping important traits as fruit colour, antioxidant activity, etc.

**Figure 1:**
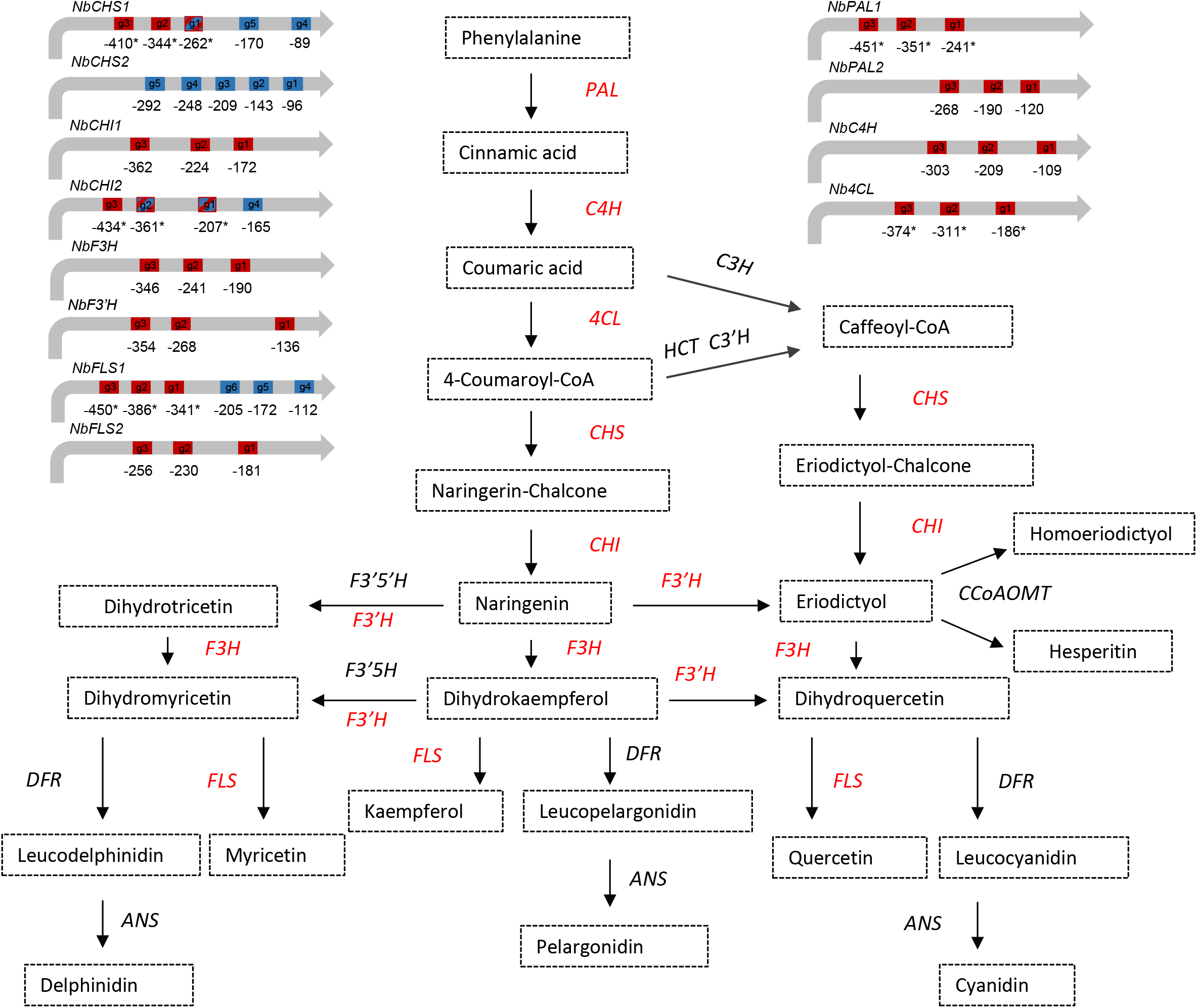
Schematic representation of the flavonoid biosynthetic pathway in plants. The different metabolites are represented in boxes. The genes that integrate the pathway are: *ANS* (anthocyanidin synthase), *CHS* (Chalcone synthase), *CHI* (Chalcone isomerase), *CCoAOMT* (Caffeoyl-CoA O-methyltransferase), *C3H* (4-coumarate 3-hydroxylase), *C3’H* (4-coumaroyl shikimate/quinate 3’-hydroxylase), *C4H* (Cinnamate 4-hydroxylase), *4CL* (4-coumaroyl CoA ligase), *DFR* (Dihydroflavonol 4-reductase), *F3H* (Flavanone 3-hydroxylase), *F3’H* (flavonoid 3’-hydroxylase), *F3’-5H* (Flavonoid 3’ 5-hydroxylase), *F3’-5’H* (Flavonoid 3’ 5’-hydroxylase), *FLS* (Flavonol synthase), *HCT* (4-hydroxycinnamoyl CoA:shikimate/quinate hydroxycinnamoyltransferase), *PAL* (Phenylalanine ammonia-lyase). The genes marked in red are those selected for transcriptional activation with dCasEV2.1 in this work. A representation of each candidate gene promoter with the gRNA positions of the first round of optimization in red and the second in blue is included. The asterisks represent the gRNA position recalculated with updated information of the TSS available in databases (https://www.nbenth.com/).

The flavonoid pathway has been the subject of remarkable metabolic engineering interventions mainly by making use of native transcription factors (Dias and Grotewold 2003; Park et al. 2021). The enzymes in the pathway are frequently regulated by a triad of TFs comprising a MYB TF, a bHLH and a WD repeat component (Zhao et al. 2013). In many cases, the overexpression of the MYB factor alone or in combination with a bHLH is sufficient to ectopically activate an entire branch of the pathway (Liu et al. 2015). MYB factors show certain degree of specificity for activating the biosynthesis of different flavonoid subgroups. Thus, whereas the *ROSEA* transcription factor activates enzymes in the anthocyanin branch in tomato and tobacco (Fresquet-Corrales et al. 2017; Vu and Lee 2019), the *Arabidopsis thaliana MYB12* factor activates the enzymes of the flavonol subgroup, leading to the accumulation of kaempferol and rutin (quercetin glycosylate) as main products (Misra et al. 2010; Zhang et al. 2015). With the elucidation of the specificities of natural TFs from different species, followed by their recombinant expression, Butelli and co-workers obtained multi-level engineering of flavonoid compounds in tomato (Butelli et al. 2008). As shown in their work, the engineering precision obtained with native TFs reaches its limit at the “subgroup” level, as native TFs collectively activate those endogenous genes sharing similar regulatory sites in their 5’untranscribed regions, and these genes usually correspond to enzymes in the same branch of the pathway (e.g., flavanols, anthocyanins or flavanones branches). To achieve higher precision levels in plant metabolic engineering, up to the level of individual enzymes (and metabolites), endogenous TF seem not to be fit to the purpose and it would be necessary to break the evolutionary constrains and incorporate the type of single-gene specificity offered by PTAs.

In this work, we aimed to explore the efficiency and the precision limits of dCas9-PTAs for engineering the specialized metabolism in plants. Using the dCasEV2.1 programmable activator previously developed in our group (Selma et al. 2019), we first selected individual activation programs (i.e. single polycistronic constructs expressing of up to six gRNAs) for ten different target enzymes distributed in the general flavonoid pathway and the flavonone/ flavonol branch of the pathway. Then, we combined those enzymes in four groups, each group leading to the biosynthesis of a different flavonoid compound as final product. Four multigene activation programs (i.e., combinations of polycistronic gRNA constructs targeting several genes simultaneously) were constructed and assayed transiently in *N. benthamiana*, each program designed to specifically activate the genes in one of the four groups. As a result, four different and highly specific metabolic profiles were generated in the leaf, with a highly predominant accumulation of the expected target products in each of the assayed combinations. These results show that dCasEV2.1 raises metabolic engineering to a new precision level, opening the door for true customization of plant metabolic composition.

## Results

### Optimization of activation programs for individual genes in the flavonoid pathway

To engineer the flavonoid biosynthesis through the custom upregulation of endogenous genes, the first step consisted in the identification of flavonoid biosynthetic genes in *Nicotiana benthamiana*, including also those encoding upstream enzymes belonging to the general phenylpropanoid pathway. The KEGG reference database with the complete flavonoid pathway was used as a guide for the selection of all candidate genes (https://www.genome.jp/kegg/pathway.html). A schematic representation of the pathway can be found in Figure 1, where the enzymatic steps intended for transcriptional activation in this work are highlighted. Candidate gene identification in the *N. benthamiana* genome was carried out manually by homology search using orthologous proteins from *A. thaliana*, *Solanum lycopersicum* and *Nicotiana tabacum* available in Uniprot (https://www.uniprot.org/) and Solgenomics database (https://solgenomics.net/). The retrieved candidates were contrasted with the automatic annotation of the version v3.3 of the *N. benthamiana* genome assembly (https://www.nbenth.com/). The allotetraploid nature of *N. benthamiana* results in several paralogs and homoeologues for each enzymatic step in the pathway, therefore the selection of candidates for transcriptional activation was performed according to the following criteria: (i) maximum homology levels with the reference proteins; (ii) completeness of gene annotation, with reliable identification of the transcriptional start site (TSS), a critical parameter that determines the region in which activation efficiency is maximum, usually between nucleotides −100 to −300 from it. (iii) optimal sequence features for gRNA design in the activation region, with an absence of putative off-targets in the *N. benthamiana* genome. On some occasions, discrepancies between *in silico* gene annotation and transcriptomic information were found. In these cases, transcriptomic data was prioritized. In total, twelve different candidate genes were selected for upregulation in two optimization rounds (see below), covering eight enzymatic steps, namely PAL, CHS, CHI, C4H, 4CL, F3’H and F3H and FLS.

Once the candidate genes were selected, their transcriptional activation was assayed individually in *N. benthamiana* transient assays employing the dCasEV2.1 activation tool. The dCasEV2.1 comprises two modules, a constant module with two constitutively expressed transcriptional units (TUs), and a variable module carrying the gene-specific activation program. In the constant module, the first TU produces a deactivated Cas9 translationally fused to an EDLL activation domain (Tiwari et al. 2012), and the second TU expresses a VPR (Chavez et al. 2015) activation domain fused to the MS2 RNA aptamer-binding protein (Konermann et al. 2015) (Figure 2A). In each assay, the activation tool was completed with the co-transformation of the target-specific module, which consists of one or two polycistronic gRNAs carrying three target-specific protospacers plus a MS2-binding RNA aptamer separated by processable tRNA spacers (Figure 2B). All protospacers were designed against the 5’untranscribed region of genes between −100 and −300 bp from the TSS. The list of the gRNAs target sites designed for each gene are listed in Table 1 and depicted in Figure 1. Four days post infiltration (dpi) the samples were collected, and levels of transcriptional activation were evaluated by RT-qPCR. The reference sample used as negative control was also infiltrated with dCasEV2.1 loaded with a non-target gRNA of *N. benthamiana* genome for a best comparison.

**Figure 2:**
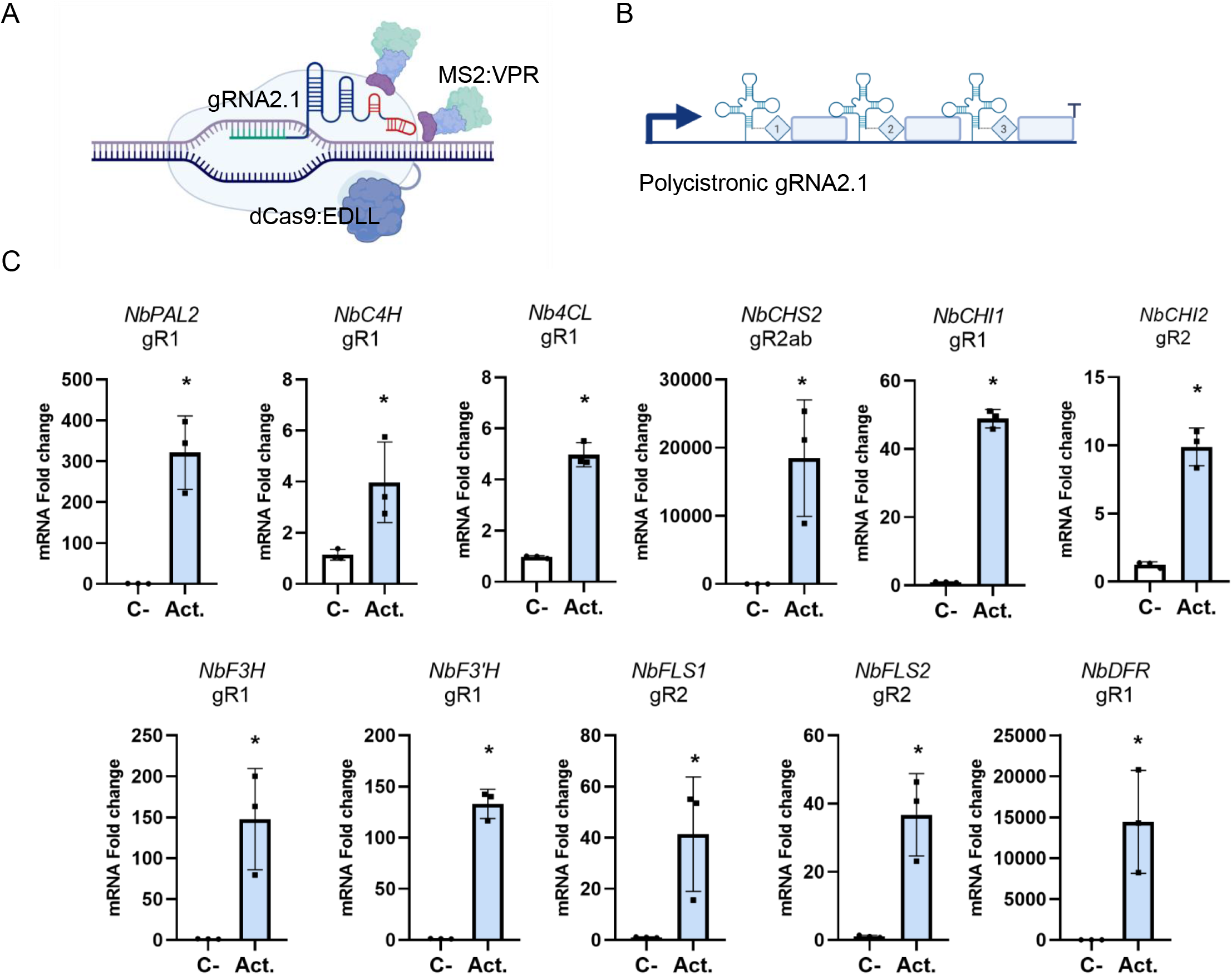
dCasEV2.1-mediated transcriptional activation of individual genes of the flavonoid pathway in *N. benthamiana*. A) Schematic representation of the dCasEV2.1 activator, comprising the dCas9 fused to the EDLL activation domain in C-terminus and the coat protein of the phage MS2 fused to the activation domain VPR. B) Schematic representation of the polycistronic gRNA2.1 array including tRNA-spaced gRNAs with MS2-binding RNA aptamers in the 3’ end of the scaffold. C) mRNA fold change at 4 dpi obtained by targeting the endogenous genes with optimized gRNAs in *N. benthamiana* leaves. The gR1 or gR2 indicate the round of optimization where the gRNAs were selected. The *NbCHS2* gR2ab activation was performed with 2 set of gRNAs. The *NbDFR* gR1 is included as internal control of activation. Bars represent average fold change +/− SD (n = 3). Asterisks indicate T student significant values (* = p<0.05). Images were created with BioRender.com.

A first round of optimization was carried out with ten selected genes, namely *NbPAL1, NbPAL2, NbC4H, NbCL4, NbCHS1, NbCHI2, NbF3H, NbF3’H*, and *NbFLS1*. The *NbDFR* gene, previously assayed in our group was also added to the analysis for comparison. Most endogenous target genes showed remarkable upregulation upon dCasEV2.1 activation treatment (see Supplementary figure 1A). Surprisingly, the activation program designed for *NbPAL1* resulted in a modest four-fold upregulation, while its homeologue *NbPAL2* showed a 200-fold activation. This was a consequence of an erroneous identification of the TSS in *NbPAL1*, which generated the design of the gRNAs in sub-optimal positions. Consequently, *NbPAL2* was selected in this work for further attempts to activate the phenylalanine ammonia-lyase. Unfortunately, the genes *NbCHS1*, *NbCHI2* and *NbFLS1* showed low activation rates in the first set of experiments. For this reason, a new optimization round for those suboptimal enzymatic steps was carried out by introducing new gRNAs design, or by targeting new alternative genes catalysing the same enzymatic step (Supplementary figure 1B). As presented in Figure 2C, after the second optimization process, a successful transcriptional activation (> 10-fold) was obtained for all selected enzymatic steps except for *NbC4H* and *Nb4CL*, where only modest four-fold activation rates were obtained. The best activation results were achieved for the previously described *NbDFR* (15000-fold) and *NbCHS2* (18000-fold).

### Programming naringenin accumulation with dCasEV2.1

The results obtained with the individual activation programs prompted us to undertake the simultaneous activation of several genes in the pathway following a modular polycistronic gRNA strategy (Lowder et al. 2017; Hashimoto et al. 2018). As a first step, we aimed at the co-activation of the enzymatic steps leading to the accumulation of naringenin, therefore involving the genes *NbPAL2*, *NbC4H*, *Nb4CL*, *NbCHS2*, *NbCHI1* and *NbCHI2*. To select the best dCasEV2.1 activation program possible, we assayed seven different gRNA multiplexing arrangements targeting all the above-mentioned genes except *NbC4H*. Each combination comprised between three to six U6-26-driven polycistronic TUs (Figure 3A). *NbC4H* was omitted as a target gene in all gRNA combinations due to its modest activation rates. All seven combined activation programs were transiently assayed in *N. benthamiana* leaves following the same methodology described for previous experiments. The RT-qPCR results in Figure 3B show that all the assayed programs resulted in significant gene activation, although induction levels were notably reduced compared with programs addressing individual genes. In general, it was observed that smaller multiplexing arrays resulted in higher activation rates of individual genes. For instance, *NbCHS2* activation reached between 800- and 1200-fold with activation programs AP-N1, AP-N1B, AP-N2 and AP-N3 comprising 2 or 3 target genes but dropped below 600-fold in programs targeting four or five genes (AP-N2B and AP-N4B). Following the same trend, those targets showing modest activations in single-gene programs showed even lower inductions with complex activation programs, dropping below significance levels in some cases such as *NbCHI2* when treated with AP-N2B or AP-N4B.

**Figure 3:**
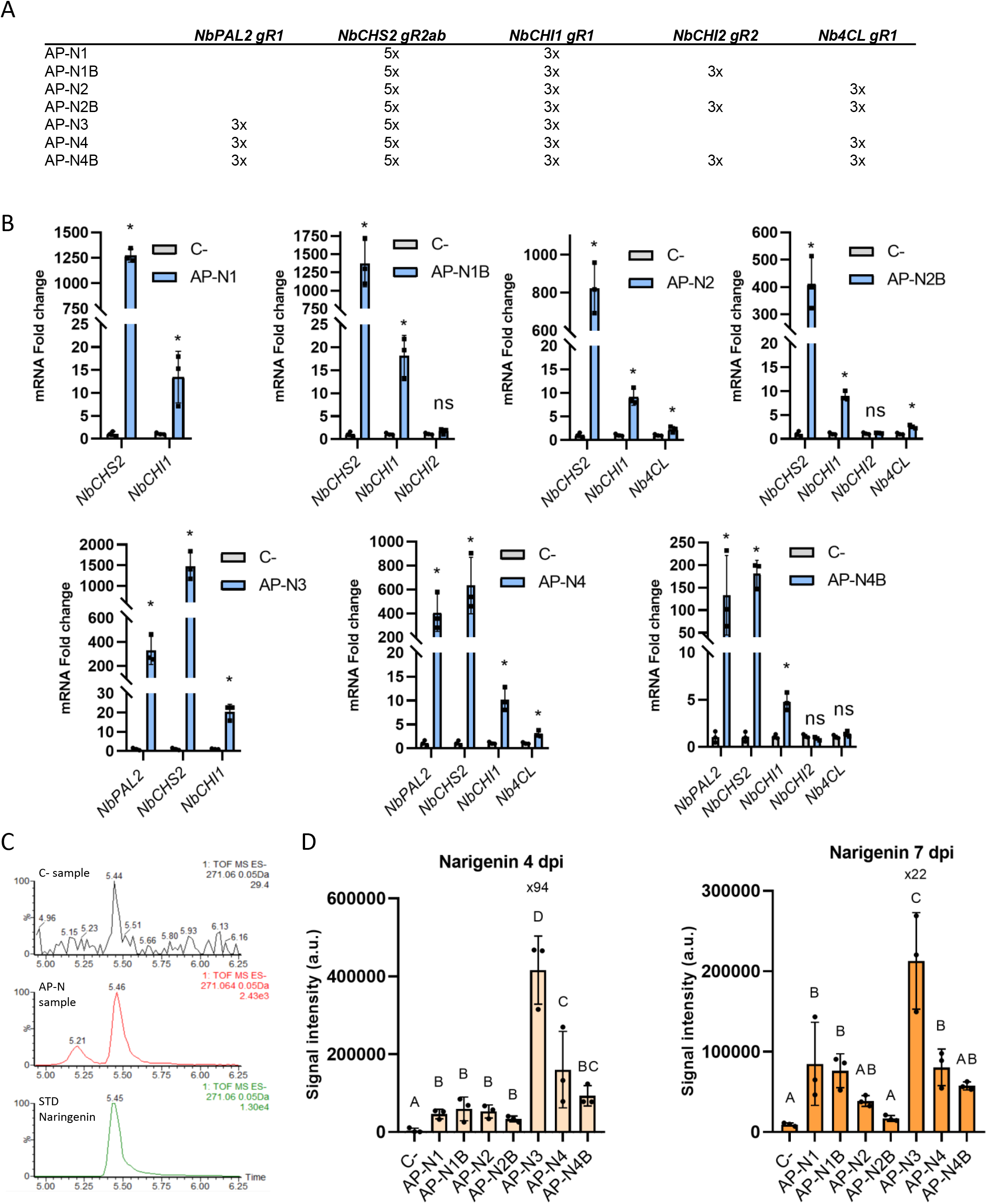
Optimization of naringenin production in *N. benthamiana* leaves through dCasEV2.1 activation. A) Representation of the different activation programs (AP)s tested for activating of naringenin production (AP-N1, AP-N1B, AP-N2, AP-N2B, AP-N3, AP-N4 and AP-N4B). B) mRNA fold change at 4 dpi obtained by targeting the endogenous genes of the flavonoid pathway with optimized gRNAs in *N. benthamiana* leaves. Bars represent average fold change +/− SD (n = 3). C) Identification of naringenin in *N. benthamiana* leaves by comparison with a true naringenin commercial standard (STD naringenin) using the ion with *m/z* 271.06 in ESI negative mode. C-sample: negative control, AP-N sample, dCasEV2.1-activated sample. D) Relative naringenin quantification in the indicated samples at 4- and 7-dpi. Bars represent average intensity. Different letters indicate statistically significant differences (ANOVA, Tukey HSD test; p<0.01, n=3). Asterisks indicate T student significant values (* = p<0.05).

In order to see if the changes in transcript profiles have resulted in the expected metabolic changes, naringenin levels in treated samples were analysed by LC-MS (UPLC-(ESI)-QTOFMS) at 4 and 7 dpi using a purified commercial standard for identification (Figure 3C). As shown in Figure 3D, naringenin content was enriched in all samples as compared with a control leaf where dCasEV2.1 was loaded with an unrelated program. Maximum levels of naringenin were obtained with AP-N3, which targeted *NbPAL2, NbCHS2* and *NbCHI1* simultaneously. In this combination, the levels of the target compound were raised almost 100-fold as compared with the levels in control samples. The upregulation of *NbPAL* seemed important for the early activation of the pathway, since the best levels of naringenin accumulation were found in samples where this enzyme was upregulated. The accumulation trends of the targeted metabolite were similar at 4 and 7 dpi, although a drop in signal intensity was observed in the later time point, particularly in samples AP-N3, AP-N4 and AP-N4B.

### Customization of flavonoid composition as a study case for single metabolite precision level engineering

Despite the indications that complex gRNA programs resulted in lower activation efficiencies than simpler ones, the remarkable levels of naringenin accumulation obtained with shorter programs suggested that there was still room for adding new instructions to AP-N3, thus extending regulation further downstream in the pathway. Furthermore, naringenin constitutes a crossroad point from which several compounds can be derived depending on the set of downstream enzymes to be activated. Thus, the activation of *F3’H* on top of AP-N3 would convert naringenin into a different flavanone, eriodictyol. Moreover, naringenin can be used as starting point for the accumulation of two important flavonols, kaempferol and quercetin. Steering the metabolic flux to produce kaempferol would require the simultaneous activation of two enzymes, first *F3H* to produce dihydrokaempferol, and next *FLS* to introduce a double bound bond in the C ring, yielding kaempferol. Alternatively, the production of quercetin can be induced by taking the eriodictyol program as a basis, and adding activation instructions for *F3H* and *FLS*, producing first dihydroquercetin and finally quercetin. Following this rationale, three new metabolite-specific gRNA programs (Figure 4A), namely AP-K, AP-E and AP-Q, were constructed and assayed in *N. benthamiana* next to AP-N3 to produce kaempferol, eriodictyol, quercetin and naringenin, respectively. Figure 4B shows the gene activation profiles obtained for each combination, where it can be observed that an increase in program complexity has a negative effect on the overall induction levels compared to simpler programs. This was clearly observed in the activation levels of *NbCHS2*, which drops by approximately 5-fold when more than four genes are targeted simultaneously. Despite the progressive reduction in activation levels, a significant upregulation is still observed in all enzymes assayed even in the more complex program (AP-K and AP-Q). It is worth noticing that *NbF3’H* is only activated in AP-Q but not in AP-K, this serving as an additional indication of the specificity of the activation programs and discarding positive feedback as cause for the observed upregulations.

**Figure 4:**
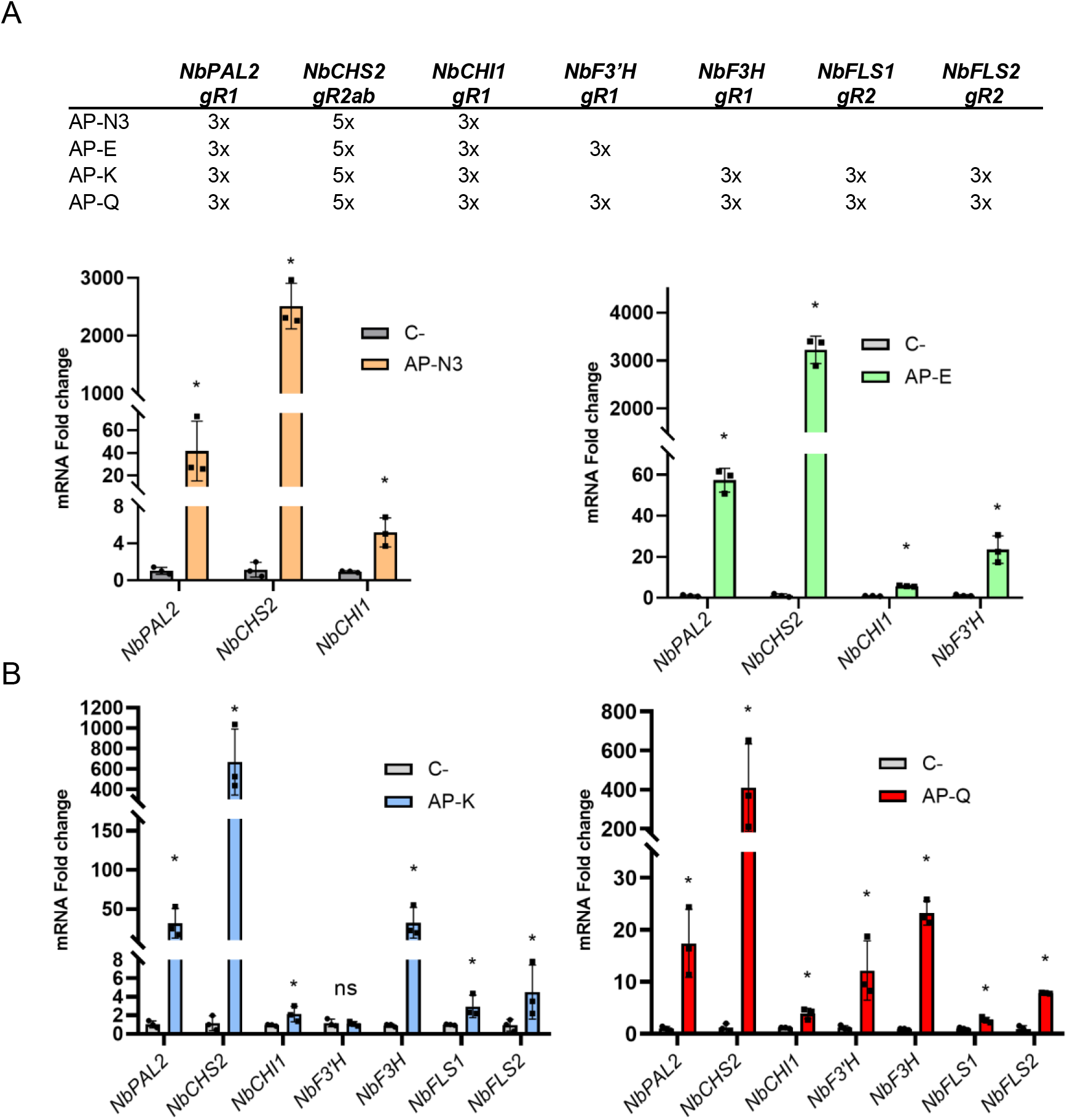
Design and relative expression analyses of activation programs (APs) for naringenin, eriodictyol, kaempferol and quercetin production in *N. benthamiana*. A) Representation of the Activation Programs (APs), the gRNAs and the targeted genes included in each of them (AP-N3 for Naringenin Activation Program, AP-E for Eriodictyol Activation Program, AP-K for Kaempferol Activation Program and AP-Q for Quercetin Activation Program). B) mRNA fold change at 5 dpi obtained by targeting the endogenous genes with optimized gRNAs in *N. benthamiana* leaves for the groups AP-N3, AP-E, AP-K and AP-Q. Bars represent average fold change +/− SD (n = 3). Asterisks indicate T student significant values (* = p<0.05)

To understand to what extent the customized activation of different subsections of the pathway led to differential flavonoid composition in treated leaves, an untargeted LC-MS metabolomic analysis was carried out with the same samples previously analysed by RT-qPCR. As the Principal Component Analysis (PCA) in Figure 5A indicates, each activation program produced a distinct and characteristic metabolite profile, with a first main component separating controls from flavonoid-activated samples accounting for 28.9% of the variance, and a second principal component separating flavonones Activation Programs to the flavonols Activation Programs accounting for 23.4% of the variance. The level of precision achieved with the four activation programs became even more evident when the 100 most significantly different features were hierarchically clustered (Figure 5B). Here, a perfect clustering was observed that parallels the activation programs and the control sample. A detailed version of the heatmap can be found in Supplementary figure 3. The differential *m/z* ions and their respective retention times are listed in Supplementary table 2. Furthermore, when metabolites in each cluster were tentatively identified attending to their retention time and the characteristic *m/z* ratios (see Materials and Methods), a remarkable match was found between the activation program employed and the predominant metabolites in the samples. Thus, as expected, AP-N3 samples accumulate naringenin aglycon and three other glycoside derivatives more than any other sample. Eriodictyol and its sugar conjugates are the metabolites accounting for the aggrupation of the APE hierarchical cluster, although certain levels of eriodictyol are also found in the AP-N3 cluster. This is not surprising since both flavanones are only one enzymatic step away from each other, and eriodictyol can be also synthetized from coumaric acid following a secondary branch in the pathway (see Figure 1). Remarkably, flavonols are almost completely absent both in AP-N3 and AP-E. On the contrary, kaempferol and its glycosylated derivatives are the main differentially accumulated compounds in AP-K samples, and conversely, the quercetin aglycon and its conjugated derivatives are most abundant in AP-Q samples. As could be anticipated, a certain level of cross-contamination is observed in both flavonol programs. Again, this is not entirely unexpected as both AP-K and AP-Q share 5 out of 6 steps in their respective programs. It is worth noting that flavonol levels remain low in AP-K and AP-Q samples, indicating that successful activation of downstream genes is responsible for the specific accumulation of flavonol compounds.

**Figure 5:**
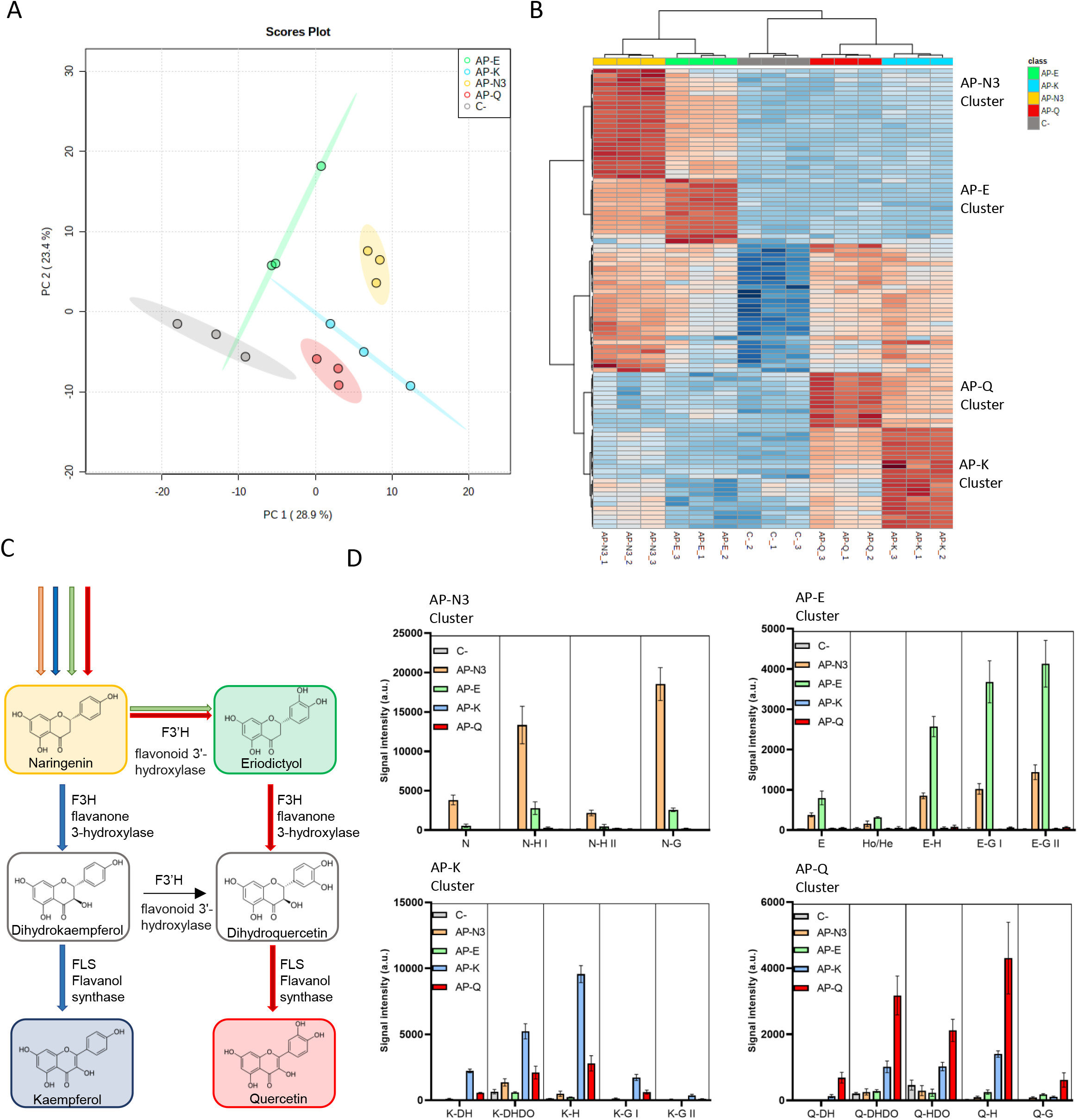
Analysis of the metabolic profiles of *N. benthamiana* leaves treated with activation programs (APs) for naringenin, eriodictyol, kaempferol and quercetin. A) Principal Component Analysis resulting from the untargeted LC-MS data obtained in ESI negative mode from leaves treated with programs AP-N3, AP-E, AP-K and AP-Q and control samples (C-) agroinfiltrated with unprogrammed dCasEV2.1. B) Hierarchical cluster analysis and heatmap representation of the Control, AP-N3, AP-E, AP-K and AP-Q metabolic profiles, with three biological replicates per condition (ESI-). The metabolites represented are the 100 most significant using a t-test analysis (p < 0.05) of each sample. The data was obtained using Euclidean distance and Ward’s minimum variance method. Red indicates up-regulated and blue downregulated features. Each *m/z* corresponds to the same compound in all samples. C) Schematic representation of selected target metabolites of the flavonoid pathway and the genes involved in each enzymatic reaction required for their biosynthesis. The coloured bars represent the targeted gene for each group (AP-N3 in orange, AP-E in green, AP-K in blue and AP-Q in red). D) Relative abundance of identified metabolites. The ions employed for the metabolic quantification are the parental ions identified in the heatmap as single compounds (See also Supplementary table 2). N, Naringenin; N-H, Naringenin Hexose; N-G. Naringenin Glucoside; E, Eriodictyol; E-H, Eriodictyol Hexose; E-G, Eriodictyol Glucoside; K-DH, Kaempferol Dihexose; K-DHDO, Kaempferol Dihexose-deoxyhexose; K-H, Kaempferol Hexose; K-G, Kaempferol Glucoside; Q-DH, Quercetin Dihexose; Q-DHDO, Dihexose-deoxyhexose; Q-HDO, Quercetin Hexose-deoxyhexose; Q-H, Quercetin Hexose; Q-G, Quercetin Glucoside.

## Discussion

Programmable transcriptional activators (PTAs) based on CRISPR/dCas9 architecture are powerful tools for the transcriptional regulation of a wide spectrum of targets. After a first generation of PTAs based on the direct translational fusion of ADs to dCas9 showed modest induction activities (Piatek et al. 2015), subsequent generations have emerged incorporating multiple AD anchoring sites such as multi-epitope chains (Papikian et al. 2019), RNA aptamers or combinations of them. These upgraded versions considerably boosted PTAs ability to produce strong activation of targeted genes (Lowder et al. 2018; Lee et al. 2019). These improvements, added to the multiplexing capacity of CRISPR/dCas9, have turned CRISPR-PTAs into extraordinary tools for Plant Synthetic Biology applications. Of particular interest is the ability offered by CRISPR-PTAs to tinkering with enzyme expression in metabolic pathways, thus shaping the plant metabolite composition. In this work we tinkered with a section of the flavonoid pathway, exploring the limits of the tool, to find out that dCasEV2.1, the CRISPR-PTA developed in our laboratory, offers an unprecedented level of precision in plant metabolic engineering interventions. We anticipate that other CRISPR-PTAs developed by other groups, having an equivalent mode of action and showing similar gene activation levels, would perform similarly well when used for the same purposes. We show that with a careful selection of gRNAs, simultaneous regulation of up to six enzymatic steps can be achieved, leading to the highly preferential accumulation of individual flavanones (naringenin or eriodictyol) and flavonols (quercetin or kaempferol). The ability to harness a pathway towards the production of a single predominant metabolite has important implications in plant biomanufacturing and biorefinery, since the isolation of pure valuable metabolites is known to be hindered by the presence of contaminant compounds belonging to the same pathway, hence showing similar chemical properties. In the examples described here, a considerable proportion of the targeted metabolites were found in glycosylated forms. *N. benthamiana*, as many other plant species, is promiscuous in glycosyl transferase activities (Wang 2009). From a biomanufacturing point of view, glycosylated forms constitute a relatively minor problem due to the affordability of industrial glycosylases. As an alternative, glycosyl transferases could be also targeted for programmable repression, leading to the predominant accumulation of the aglycon.

In this work we only use transcriptional activators as tools to re-route metabolic fluxes. Programmable repression is partially dispensable for the control of the flavonoid pathway because in *N. benthamiana* leaves the flux through the pathway is low. However, the interplay of programmable repressors would enable to further refine the accuracy of re-routing programs. As mentioned above, the repression of glycosyl transferases would serve to ensure the predominance of aglycon forms if required. Furthermore, as shown in Figure 1, eriodictyol can be synthetized by an alternative route that involves the action of *NbC3’H, NbC3H* and *NbHCT* genes. Programmed repression of one or more of these genes could contribute to reduce the levels of accompanying metabolites when the production of a single flavonol is attempted. Furthermore, programmable repression will be strongly required for harnessing metabolic pathways other than flavonoid pathway which are highly active by default. Unfortunately, there are few examples in the literature showing a highly efficient repression based on CRISPR architecture (Tang et al. 2017). All currently described tools, although helpful, are probably insufficient in providing full control due to their inability to conduct strong transcriptional repressions in highly active genes. It has been proposed that dCas9, due to its mode of action when binding DNA, which implies a relaxation of the chromatin in the surrounding area, has limited capacity to act as a strong repressor when fused to conventional repressor domains. This limitation might be circumvented by the use of epigenetic repressors adding reversible chromatin silencing marks (Gallego-Bartolomé 2020). Alternatively, programmable repression could be achieved by other means, such as post-transcriptional gene silencing tools (Mahas et al. 2019). Given the remarkable ability of dCasEV2.1 to program transcriptional activation, its combination with efficient repressors would enable near full control of metabolic pathways in plants.

For the delivery of activation programs, we made use of transient *Agrobacterium*-based transformation. Transient delivery of genetic information in the form of T-DNA or RNA is becoming increasingly popular in plant biotechnology. Recently, transient reprogramming of crop plants was shown using RNA virus-based delivery systems, either as viral particles or mediated by *Agrobacterium* (Torti et al. 2021). Agroinfiltration has become not only a widely used experimental procedure (Norkunas et al. 2018; Grosse-Holz et al. 2018), but also a potent and scalable plant biomanufacturing strategy as recently demonstrated with the production of plant-made vaccines against influenza and SARS-CoV2 (D’Aoust et al. 2008; Diego-Martin et al. 2020). On the other hand, the gRNA elements in PTAs can be also transiently delivered using viral vectors, as recently shown using TRV as delivery agent (Ghoshal et al. 2020). Reprogramming metabolic pathways using transient tools has the additional advantage of circumventing the need for stable transgenics, giving regulatory advantages in some areas. Alternatively, stable transformation of the activation programs could also be envisioned as a powerful tool in crop breeding. Stable integration in the plant’s genome could circumvent the complexity limits that have been evidenced in this work, which could be attributed to the cargo capacity of the T-DNAs or its ability to cope with highly repetitive gRNA structures. Stable transgenic programs could be pyramided in different genomic loci, avoiding limits imposed by T-DNA cargo capacity. If required by the presence of many gRNA-coded instructions, the expression of the remaining components in dCasEV2.1 could be boosted with state-of-the-art strategies, preventing them from becoming a limiting factor for activation (Pasin et al. 2017). Connecting integrated programs with endogenous or exogenous cues using appropriated genetic sensors (Ochoa-Fernandez et al. 2020; Randall 2021) would provide the ultimate ability to customize plant metabolic composition using its own endogenous metabolic pathways.

In sum, we show here that CRISPR/dCas9-based transcriptional activators provide the sufficient precision and multiplexing capacity to reprogram metabolic pathways and customize metabolic composition in plants. This ability has important implications in feed and food design, as well as in the valorisation of industrial crops.

## Methods

### DNA constructs

All plasmids used in this work were assembled using GoldenBraid (GB) cloning (Sarrion-Perdigones et al. 2013). The DNA sequences of the constructs generated in this work are available at https://gbcloning.upv.es/ by entering the IDs provided in Supplementary table 3. Briefly, the multiplexing vectors used for this work were generated as GB level −1 vectors and previously described in Selma et al. (Selma et al. 2019). The level −1 vectors, pVD1_M1-3pTRNA scf 2.1 (GB1436) pVD1_M2-3pTRNA scf 2.1 (GB1437) and pVD1_M3-3pTRNA scf 2.1 (GB1438) were used to assemble the protospacer sequences occupying the first, second and third positions in the final gRNA assembly. For GB gRNA assemblies two partially complementary primers containing the protospacer sequence were designed at https://gbcloning.upv.es/do/crispr/multi_cas9_gRNA_domesticator_1 by entering the protospacer sequences listed in Supplementary table 1. Primers were resuspended in water to final concentrations of 1 μM and equal volumes of forward and reverse primer were mixed. The mixture was incubated at room temperature for 5 min for the hybridization of the primer pair. 1 μl of the primer mix was included in a BsmBI restriction–ligation reaction with 75 ng of pUPD2 and 75 ng of the corresponding level −1 vector for the assembly of tRNA-protospacer-scaffold units in level 0. The next step consists in the assembly of the multiplexing gRNA expression cassette in level 1. For level 1 assemblies, 75 ng of level 0 gRNA for each position, 75 ng of U626 promoter (GB1001), and 75 ng of pDGB3α destination vector were included in a BsaI restriction-ligation reaction.

The combination of level 1 multiplexing gRNAs and dCasEV2.1 TUs were performed by binary BsaI or BsmBI restriction–ligation reactions to obtain all the level ≥1 assemblies as it was described in Sarrion-Perdigones, A. et al. (Sarrion-Perdigones et al. 2013)

### Nicotiana benthamiana agroinfiltration

The transient expression assays were carried out through agroinfiltration *of N. benthamiana* leaves. *N. benthamiana* plants were grown for 5 weeks before agroinfiltration in a growing chamber where the growing conditions were 24 °C/20 °C light/darkness with a 16 h/8 h photoperiod. The plasmids were transferred to *A. tumefaciens* strain GV3101 by electroporation. Agroinfiltration was carried out with overnight grown bacterial cultures. The cultures were pelleted and resuspended in agroinfiltration solution (10 mM MES, pH 5.6, 10 mM MgCl2, and 200 μM acetosyringone). After incubation for 2 h at room temperature with agitation, the optical density of the bacterial cultures was adjusted to 0.1 at 600nm and mixed for co-infiltration with the silencing suppressor P19. Agroinfiltrations were carried out through the abaxial surface of the three youngest fully expanded leaves of each plant with a 1 ml needle-free syringe.

### RNA isolation and RT-qPCR Gene Expression Analysis

Total RNA was isolated from 100 mg agroinfiltrated leaves using the Gene Jet Plant Purification Mini Kit (Thermo Fisher Scientific) according to the manufacturer’s instructions. The timing for sample collection was 4 days post infiltration (dpi) for testing individual genes activation and for naringenin optimization and 5 dpi in the case of AP-N3, AP-E, AP-K, AP-Q activation. Before cDNA synthesis, total RNA was treated with the rDNAse-I Invitrogen Kit according to the manufacturer instructions. An aliquot of 1 μg of DNAse treated RNA was used for cDNA synthesis using the PrimeScript™ RT-PCR Kit (Takara) in a final volume of 20 μL according to the manufacturer indications. Expression levels for each gene were measured in triplicated reactions, in the presence of a fluorescent dye (SYBR® Premix Ex Taq) using Applied biosystem 7500 Fast Real Time PCR system with specific primer pairs (Supplementary table 4). The F-box gene was used as internal reference (Liu et al. 2012). Basal expression levels were calculated with samples agroinfiltrated with dCasEV2.1 in combination with an unspecific gRNA. mRNA fold change calculations for each sample were carried out according the comparative ΔΔCT method (Livak and Schmittgen 2001).

### Liquid chromatography (LC) and ESI mass spectrometry (MS) for flavonoids content analysis

Leaf samples of three different plants agroinfiltrated with each construct were collected at 4 and 5 dpi respectively and used as triplicates for metabolomics analyses. The same tissue was used for transcriptomics and metabolomics analyses. The tissue was frozen in liquid N_2_ and powdered with a grind mill and, finally, lyophilized. Thirty milligrams of dried tissue were extracted at 4°C with 1 mL of methanol 30% containing 0.01% of formic acid. The preparation was homogenized with a grind mill and kept in ice during 20 min. After that, the samples were centrifugated at 15000 rpm for 15 minutes. The supernatant was collected and filtrated with 20 microns cellulose strainer (Regenerated Cellulose Filter, Teknokroma). Three independent biological and two technical replicates per sample were analysed. 20 μl of each sample were injected into an Acquity UPLC system (Waters) coupled to a hybrid quadrupole time-of-flight instrument (QTOF MS Premier). Separation was performed using an HPLC SunFire C18 analytical column with a particle size of 5 μm, 2.1 × 100 mm (Waters). A gradient of methanol and water containing 0.01% formic acid was used. The gradient started with 95% aqueous solvent and a flow of 0.3 ml per minute. The gradient reached 50% of aqueous solvent at 8 min, increasing the level of organic solvent to 95% at 12 minutes. The gradient was kept in isocratic conditions for 1 min and later returned to initial conditions in 2 minutes. The column could equilibrate for 3 minutes, for a total of 22 minutes per sample.

Data were collected in MS and MS/MS mode to gain structural information of the detected metabolites. The MS/MS function was programmed in a range of 5 to 45 eV t-wave to obtain each analyte fragmentation spectrum (Pastor et al. 2018). The electrospray ionization was performed in positive and negative mode and analysed individually in order to obtain a best profile of the flavonoid metabolites following the specifications described by Gamir et al. (Gamir et al. 2012).

For unequivocal metabolite determination, samples of naringenin and eriodictyol chemical standard (Sigma) were analysed in the same conditions. The exact mass, specific retention time, and spectrum fragmentation of naringenin and eriodyctiol standard were compared to the fragmentation profiles of each sample as described by Schymanski et al. (Schymanski et al. 2014).

### Naringenin content data analysis and statistics

The raw data obtained were processed by Masslynx 4.1 software and transformed to .cdf files. The positive (ESI+) and negative (ESI-) signals were analysed separately. Naringenin identification was carried out by comparison with a purified standard also analysed in the same conditions.

The quasimolecular ion with *m/z* 271.06 in ESI negative mode, retention time (5.47 min.) and the fragmentarion ions *m/z* 151 and *m/z* 119 allowed to unequivocally identify naringenin in the samples (Supplementary figure 2).

Metabolite amounts were quantified based on the normalized peak area units relative to their respective dry weight. All statistical analyses were conducted with Statgraphics Centurion software for the ANOVA statistical analysis (p < 0.05) and means were expressed with the standard error.

### Untargeted data analysis and statistics

The raw data obtained were processed and transformed to .cdf files. The negative (ESI-) signals were analysed employing the xcms software (https://xcmsonline.scripps.edu/) for comparing all the samples. TIC normalization was applied to each biological triplicate with a baseline of 20 peak intensity relative units. All peaks with a signal lower than 300 in all samples were eliminated to reduce the background. The data analysed comprises the retention time between 1 to 6.5 min (corresponding to the elution conditions for phenolic compounds) and *m/z* values ranging 200 to 1000. Finally, a cut-off of 75% in the coefficient of variation was applied between biological triplicates.

The data obtained was analysed using the MetaboAnalyst5.0 Software (https://www.metaboanalyst.ca/). Logarithmic transformation and Pareto scaling were employed as normalization to elaborate the Principal Component Analysis and the Hierarchical Clustering and Heatmap. Euclidean distance and Ward Clustering algorithm were applied as parameters for elaborating Hierarchical Clustering, and an ANOVA test was the statistical method used for generated the 100 *m/z* more significantly different in each group.

The *m/z* values obtained as significantly different in each group were manually clustered into single metabolites employing the original chromatograms. Finally, the tentative identification of each metabolite was carried out using external databases (https://pubchem.ncbi.nlm.nih.gov/) and the information obtained with their fragmentation profiles and collected in Supplementary table 2. The quantification of the metabolites was performed employing the parental ion identified in the F1.

## Supporting information

Supplementary figures captation

## Author contributions

D.O. and S.S. designed the experiments. S.S. and N.S, conducted the experiments. A-R.A. and L-G.M contributes with data analysis. D.O. and S.S drafted the manuscript. G.S V-V.M. F.V and G.T contribute with the manuscript writing and editing. All the authors discussed and revised the manuscript.

## Acknowledgements

This work has been funded by Grant BIO2016-78601-R and PID2019-108203RB-10 Plan Nacional I+D, Spanish Ministry of Economy and Competitiveness and Spanish Ministry of Science and Innovation. Sara Selma is a recipient of FPI fellowship associated with this Grant (BES-2017-080098).

## Abbreviation list

4CL: 4-coumaroyl CoA ligase
C4H: Cinnamate 4-hydroxylase
CHI: Chalcone isomerase
CHS: Chalcone synthase
DFR: Dihydroflavonol 4-reductase
F3’H: Flavonoid 3’-hydroxylase
F3H: Flavanone 3-hydroxylase
FLS: Flavonol synthase
PAL: Phenylalanine ammonia-lyase
VPR: VP64-P65-RtA

