## Supplementary figures and images for "Custom-made design of metabolite composition in *N. benthamiana* leaves using CRISPR activators"

### Supplementary figure 1

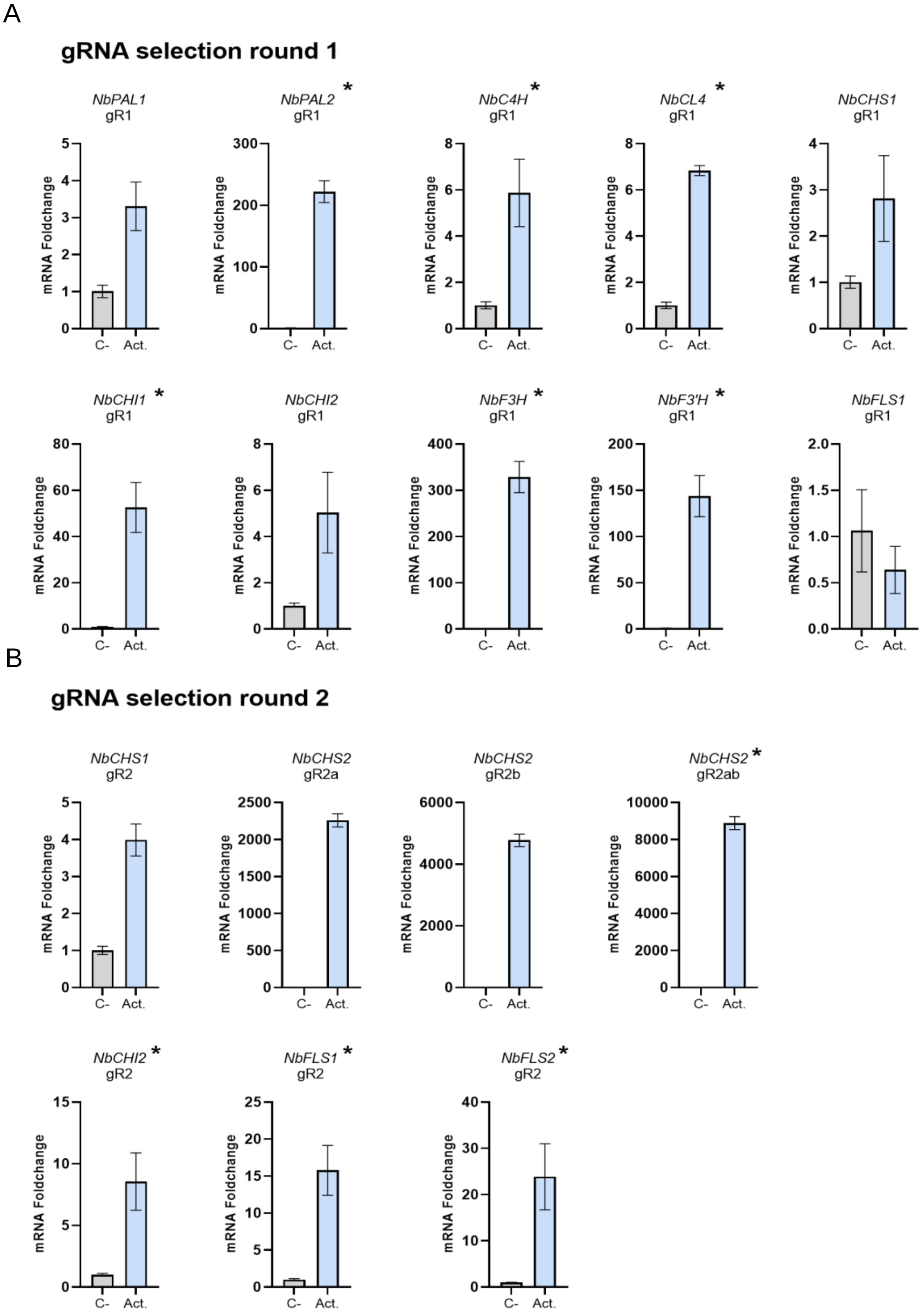

### Supplementary figure 2

Spectrum (ESI –) Naringenin m/z = 271.064    RT = 5.45 min

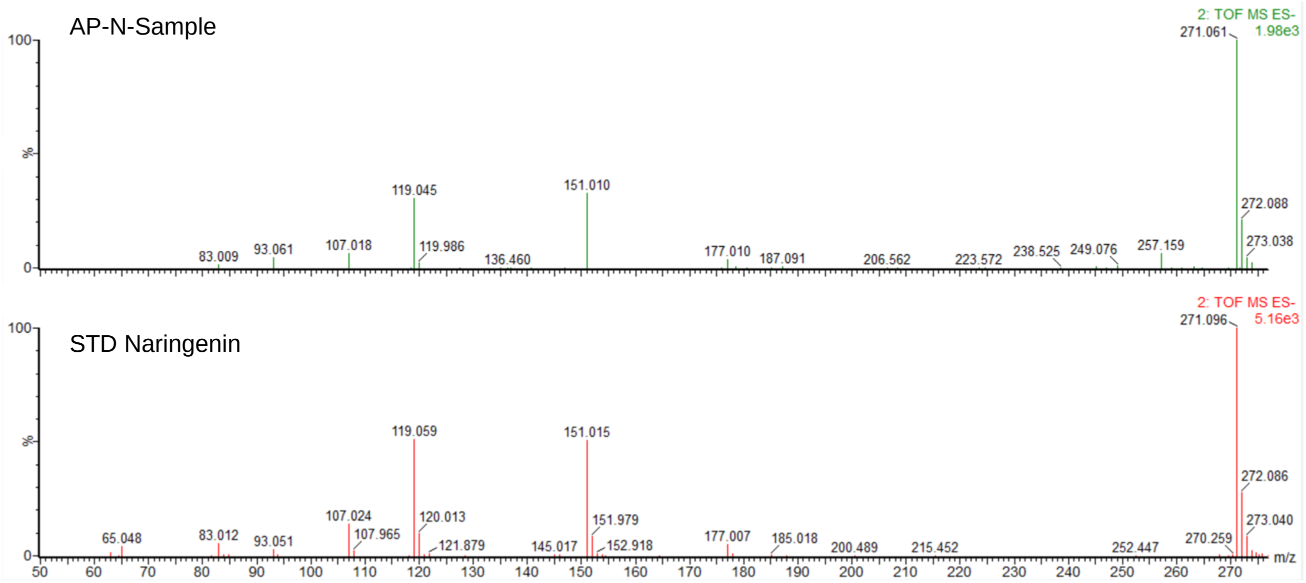

### Supplementary figure 3

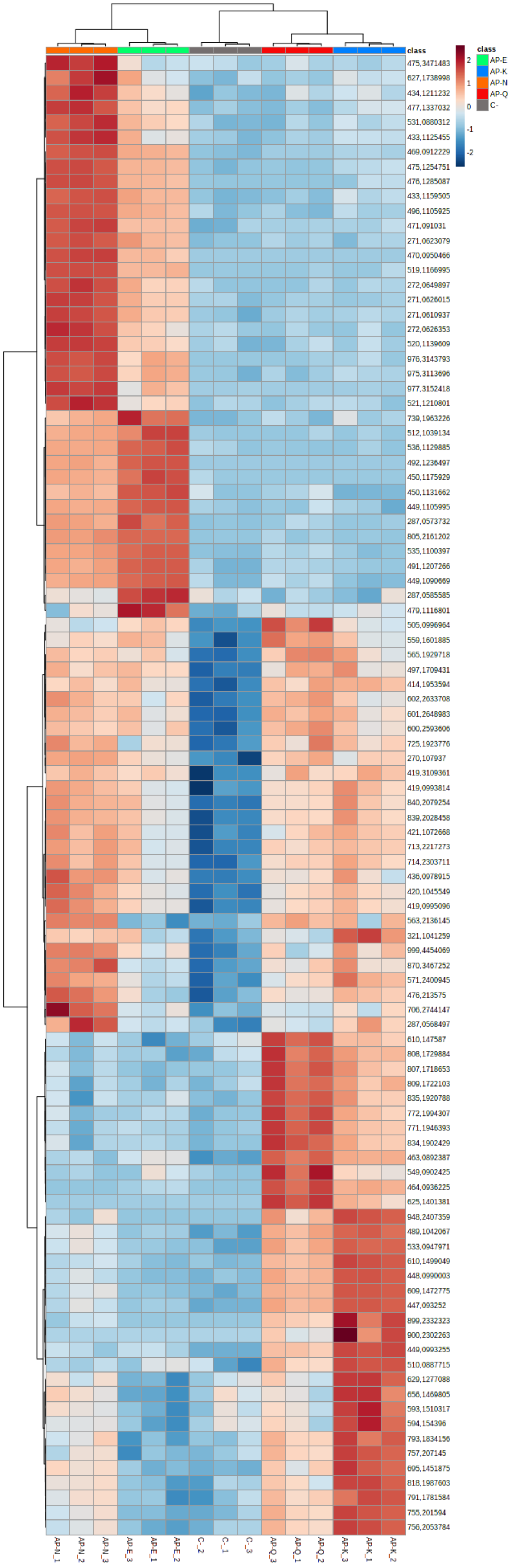
