## Supplementary figures captation for "Custom-made design of metabolite composition in *N. benthamiana* leaves using CRISPR activators"

### Supplementary figures caption:

**Supplementary figure 1: gRNA optimization and selection.** A) mRNA fold change at 4 dpi obtained by targeting the endogenous genes of flavonoid pathway with the first round of gRNA selection for each gene. B) mRNA fold change at 4 dpi obtained by targeting the endogenous genes of flavonoid pathway with the second round of gRNA selection for each gene. The asterisks represent the combination of gRNA that were selected for the following assays. Bars represent average fold change +/- SD, n = 3 technical replicates in all experiments.

**Supplementary figure 2: Negative-ion mode fragmentation of naringenin**. MS-MS spectra of AP-N sample and naringenin commercial standard (STD Naringenin).

**Supplementary figure 3: Detailed version of hierarchical cluster analysis and heatmap representation of metabolic profile of C- AP-N3, AP-E, AP-K and G AP-Q samples**. The metabolites represented are the top 100 more significant using a t-test analysis ($p < 0.05$). The data was obtained using Euclidean distance and Ward's minimum variance method. Red box indicates up-regulated compounds. Each *m/z* corresponds to the same compound in all samples.
