## Supplementary table 1 for "Custom-made design of metabolite composition in *N. benthamiana* leaves using CRISPR activators"

| Gene | Solgenomics Accesion | Nbenth.com Accesion | gRNAs sequence gR1 | gRNAs sequence gR2 |
|---|---|---|---|---|
| *NbPAL1* | Niben101Scf04656g01001 | NbLab330C12: NbE44070663.1 | gRNA1: CCTAATTAATATTCTTGAAA<br>gRNA2: CAATAATTGTTTTCAGAAAA<br>gRNA3: TCACCACTATTAATATAAGA | |
| *NbPAL2* | Niben101Scf02432g00011 | NbLab330C16: 80768655…80771248 | gRNA1: TGAAAAGCTAGACTGACCGT<br>gRNA2: GTTGAAGGGTGGTTGTTGAG<br>gRNA3: ATATTGTTTCATATTATTAC | |
| *NbCHS1* | Niben101Scf02893g00001.1 | NbLab330C10: NbD021251.1 | gRNA1: TAATTCTTAAGATTTCTAAT<br>gRNA2: TCCTAAAATTATAATGTATG<br>gRNA3: GGATTTGATTTTAAAGATAT | gRNA1: TAATTCTTAAGATTTCTAAT<br>gRNA4: GAAGTGCATTATGTTTGCAT<br>gRNA5: AGAAAATATCATTTGCAAGA |
| *NbCHS2* | Niben101scf00536g13015 | NbLab330C10: NbD004910.1 | | gR2a:<br>gRNA1: AGGTCACGTGATCTTAATGT<br>gRNA2: TGCTGAGAGAAGAGTTGAAA<br>gRNA3: TACCTACATCTCAAGATCAA<br>gR2b:<br>gRNA1: AGGTCACGTGATCTTAATGT<br>gRNA4: AATCTATTGAATCAAATATA<br>gRNA5: CCAAAGCATCATTCTTCAAA |
| *NbCHI1* | Niben101Scf05989g01008 | NbLab330C11: NbD035233.1 | gRNA1: AAAGTTGGGTGAATCCTTTA<br>gRNA2: GTCTATATAATTGTTACTTA<br>gRNA3: ATGAAGGTAAACGATAGAAT | |
| *NbCHI2* | NbLab330C10: NbD025871. | NbLab330C01: NbD014654.1 | gRNA1: ACAAGCTGTTAAGGAAAGTT<br>gRNA2: | gRNA1: ACAAGCTGTTAAGGAAAGTT<br>gRNA2: |

| | | | | |
|---|---|---|---|---|
| | 1 | | TTAAGAAAAAGTTACTCGTG<br><br>gRNA3:<br>TATCTTTTCAAAAAAAATCT | TTAAGAAAAAGTTACTCGTG<br><br>gRNA4:<br>CGAGAGAGAGGGTTGACTGG |
| *NbC4H* | Niben101Scf<br>08196g0100<br>7 | NbLab330C00:<br>NbD041626.1 | gRNA1:<br>AACTTAAGCAGAAAATGACG<br>gRNA2:<br>ATAACCTTGTGTAAATGAAA<br><br>gRNA3:<br>TTAAAGGATAAAAGCTTTAT | |
| *Nb4CL* | Niben101Scf<br>07623g0103<br>6 | NbLab330C07:<br>NbD040166.1 | gRNA1:<br>TAGACATATTAGAAGACAGG<br><br>gRNA2:<br>TATTTCTATCTTGGGTAAAG<br><br>gRNA3:<br>TGAGAGTAGTTCAGGGAATG | |
| *NbF3H* | Niben101Scf<br>09345g0000<br>4 | NbLab330C02:<br>NbD044172.1 | gRNA1:<br>TCCATTGCTTCTGAGTTGAA<br><br>gRNA2:<br>TATTTTGTTAATTAAAATTT<br><br>gRNA3:<br>ACGGAGGAATACACTATTTT | |
| *NbF3’H* | Niben101Scf<br>00974g0101<br>8 | NbLab330C06:<br>NbD007916.1 | gRNA1:<br>TTTTTTGGAATCGACTAGAA<br><br>gRNA2:<br>AAGAAGTAGATTTTCAAAAT<br><br>gRNA3:<br>AAATATTTAAAGCACGAGGT | |
| *NbFLS1* | Niben101Scf<br>02429g0600<br>7 | NbLab330C03<br>NbD018272.1 | gRNA1:<br>ATTTTGAAATAAAATATGTA<br><br>gRNA2:<br>TTAAATTTATTCTTGATGAA<br><br>gRNA3:<br>ATCATATTTCTAAAAGAACG | gRNA4:<br>GCCTATACCTAACACTCTCA<br><br>gRNA5:<br>TTTCTCGTGGGTGACTTACG<br><br>gRNA6:<br>TTACACGAACAAGTGAAGCT |
| *NbFLS2* | Niben101Scf<br>03779g0400<br>1 | NbLab330C10<br><br>NbD025871.1 | | gRNA1:<br>GTACATTCATGTGAACATGT<br><br>gRNA2:<br>GGAATGACTAGGATTGAAAA<br><br>gRNA3:<br>TCTTCATATTCGTTTTCTGT |
| *NbDFR* | Niben101Scf<br>00305g0503<br>5 | NbLab330C19:<br>NbD003043.1 | gRNA1:<br>ATGACTGACTGGTTGGTGAG<br><br>gRNA2:<br>TATCCGTATGCCTTACCTTT<br><br>gRNA3: | |

| | | | |
|---|---|---|---|
| | | | TTGAGAATTTGGTAAAACGA |

Supplementary table 1: List of candidate flavonoid genes identified in this work in *N. benthamiana*. Gene accessions for solgenomics.com and nbenth.com are provided. The protospacer sequences designed for each round of optimization are included for all tested genes.
