## Supplementary table 2 for "Custom-made design of metabolite composition in *N. benthamiana* leaves using CRISPR activators"

| Most abundat in AP: | m/z detected in F1 | Retention Time (min) | F2 (Fragmentation) of the parental ion | Tentative identification |
|---|---|---|---|---|
| AP-N3 | 469,091 | 4,31 | 469,433, 271, 193, 151 | Naringenin Hexose I |
| AP-N3 | 271,062 | | | |
| AP-N3 | 471,091 | | | |
| AP-N3 | 531,088 | | | |
| AP-N3 | 627,173 | | | |
| AP-N3 | **433,115** | | | |
| AP-N3 | 496,110 | | | |
| AP-N3 | **433,112** | 4,49 | 433, 271, 151, 109 | Naringenin Hexose II |
| AP-N3 | 271,062 | 5,15 | 475, 271, 151 | Naringenin Glucoside I |
| AP-N3 | 475,347 | | | |
| AP-N3 | 476,128 | | | |
| AP-N3 | 976,314 | | | |
| AP-N3 | 520,113 | | | |
| AP-N3 | 977,315 | | | |
| AP-N3 | 519,116 | | | |
| AP-N3 | 477,133 | | | |
| AP-N3 | 521,121 | | | |
| AP-N3 | 975,311 | | | |
| AP-N3 | **475,125** | | | |
| AP-N3 | **271,061** | 5,37 | 151, 119 | Naringenin (STD)* |
| AP-E | 287,058 | 3,90 | 739, 512,547,485 449,287,151 | Eriodictyol Glucoside I |
| AP-E | **449,109** | | | |
| AP-E | 450,113 | | | |
| AP-E | 512,103 | | | |
| AP-E | 805,216 | | | |
| AP-E | 450,113 | 4,58 | 287 | Eriodictyol Hexose |
| AP-E | **449,110** | | | |
| AP-E | 536,114 | 4,80 | 535, 491,287,151 | Eriyodictyol Glucoside II |

| | | | | |
|---|---|---|---|---|
| AP-E | **535,109** | | | |
| AP-E | 492,123 | | | |
| AP-E | 491,121 | | | |
| AP-E | 287,057 | 4,88 | 151 | Eriodictyol (STD)* |
| AP-E | 301,061 | 5,15 | 287,271,151 | Homoeriodictyol/Hespertin |
| AP-K | 900,230 | 3,67 | 609,579 447, 285 | Kaempferol Dihexose |
| AP-K | **609,147** | | | |
| AP-K | 899,233 | | | |
| AP-K | 610,149 | | | |
| AP-K | 948,240 | | | |
| AP-K | 818,198 | 3,75 | 593,449, 289,285 | Kaempferol Dihexose-dioxyhexose |
| AP-K | 793,183 | | | |
| AP-K | **755,201** | | | |
| AP-K | 756,205 | | | |
| AP-K | 757,207 | | | |
| AP-K | 791,178 | | | |
| AP-K | 695,145 | 4,3 | 465, 289,285, 245 | Kaemferol Glucoside II |
| AP-K | **447,093** | 4,93 | 447, 285,133,115 | Kaempferol Hexose |
| AP-K | 448,099 | | | |
| AP-K | 449,099 | | | |
| AP-K | 629,127 | | | |
| AP-K | 593,151 | | | |
| AP-K | 594,154 | | | |
| AP-K | 656,146 | | | |
| AP-K | 510,088 | | | |
| AP-K | 533,094 | 5,70 | 489,433,285,173 | Kaempferol Glucoside I |
| AP-K | **489,104** | | | |
| AP-Q | **625,140** | 3,38 | 609,465,301,299 | Quercetin-dihexose |
| AP-Q | 835,192 | 3,45 | 609,301 | Quercetin Dihexose- |

| | | | | |
|---|---|---|---|---|
| AP-Q | 834,190 | | | dioxyhexose |
| AP-Q | 807,171 | | | |
| AP-Q | **771,194** | | | |
| AP-Q | 772,199 | | | |
| AP-Q | 808,172 | | | |
| AP-Q | 809,172 | | | |
| AP-Q | **609,145** | 4,55 | 463, 301,300 | Quercetin Hexose-dioxyhexose |
| AP-Q | 610,147 | | | |
| AP-Q | **463,089** | 4,58 | 301,300 | QuercetinHexose |
| AP-Q | 464,093 | | | |
| AP-Q | 549,090 | 5,34 | 537,505,469,301,300 | Quercetin Glucoside I |
| AP-Q | **505,099** | | | |

Supplementary table 2: Metabolite tentative identification in AP-N, AP- E, AP- K and AP- Q samples. The *m/z* represented were identified as significant differential in the hierarchical clustering and heatmap. The m/z marked in bold represents the parental ion in negative ESI – used for quantification. The *m/z* were grouped by retention time and checked with the second fragmentation (F2) of the chromatogram for the correct identification. (STD)* represents the metabolites identified with purified standards.
