## Supplementary table 3 for "Custom-made design of metabolite composition in *N. benthamiana* leaves using CRISPR activators"

| GB_ID | Construct | GB | Construct |
|---|---|---|---|
| GB2085 | dCasEV | GB2394 | NbFLS1 gR1 |
| GB2070 | Non-target gRNA2.1 | GB2639 | NbFLS1 gR2 |
| GB2389 | NbPAL1 gR1 | GB2530 | NbFLS2 gR2 |
| GB2396 | NbPAL2 gR1 | GB2170 | NbDFR gR1 |
| GB2760 | NbCL4 gR1 | GB2641 | AP-N3 |
| GB2390 | NbCHS1 gR1 | GB2777 | AP-N1 |
| GB2500 | NbCHS2 gR2a | GB2776 | AP-N1B |
| GB2397 | NbCHS2 gR2b | GB2864 | AP-N2 |
| GB2599 | NbCHS2 gR2ab | GB2863 | AP-N2B |
| GB2502 | NbCHI1 gR1 | GB2866 | AP-N4 |
| GB2391 | NbCHI2 gR1 | GB2867 | AP-N4B |
| GB2503 | NbCHI2 gR2 | GB3259 | AP-K |
| GB2392 | NbF3H gR1 | GB3242 | AP-E |
| GB2531 | NbF3'H gR1 | GB3243 | AP-Q |
| GB2395 | NbC4H gR1 | | |

Supplementary table 3: List of GoldenBraid plasmids used in this work. DNA sequences can be found at https://gbcloning.upv.es/ by entering the provided GB_ID.
