## Supplementary table 4 for "Custom-made design of metabolite composition in *N. benthamiana* leaves using CRISPR activators"

| Gene | Primer Sequence5'-3' |
|---|---|
| *NbPAL1* | Fw: CGACAAGAGCGGCAATGCTAGTGAG<br>RV: CGGAGAGGCAAGCATGGGGTGATAT |
| *NbPAL2* | Fw: AAGATCGCAAAGCACATTTATTTG<br>RV: CGAAGTATATGTGATGTTTTGATGGA |
| *NbCHS1* | Fw: GCATTCCAACCATTAGGTCTTTCGGA<br>RV: CGAAGTTTCTCTGGCTTTAGGCTCAA |
| *NbCHS2* | Fw: ACTACTGGTGAAGGGCTTGAATGGG<br>RV: GTCCTGCCCACTAAGTAGCAACACT |
| *NbCHI1* | Fw: AATCCTATGAGACAACTGTTGGCCC<br>RV: TGCATTTCACATGCTTGAGTTGACC |
| *NbCHI2* | Fw: CCGGATAGGAATTGGCTAAGATCAT<br>RV: TCTTTTCTCCTCGAGAGTTAAGGTC |
| *NbC4H* | Fw: GTGTGGGACTAAAAGAGGGATTGCC<br>RV: GAGTAGGAGTGCAAATCACTGAGCC |
| *Nb4CL* | Fw: GGGCCATTTGTGCCGCCTATTGTTC<br>RV: GCAGCCCCAGACATGACAGTCCTTAC |
| *NbF3H* | Fw: GGATTACTGTTCAGCCCGTTGAAGG<br>RV: CTGCTGCTATTCGAGTTCACCACTG |
| *NbF3'H* | Fw: AGAGGGGTGGATGGATGAAGCCTTA<br>RV: CAAAGTGGTTGGAAAGAACCGGCTC |
| *NbDFR* | Fw: TTCATCTGCGCATCCCATCA<br>RV: TCCCTACTGAGTTTAAAGGTATCGA |
| *NbFLS1* | Fw: GCAGAAGGAGGCCAAATCTTATGGAAC<br>RV: GAGATGATTGCAAAGAATCCAGGGTCG |
| *NbFLS2* | Fw: TTAGGCCTTTCCCCACATTCGGATC<br>RV: CTGGCTTTACAGTGATCCAAACGCC |

Supplementary table 4: Primer pairs for RT-qPCR analyses.
